# LoGoPPI enables fast and accurate protein–protein interaction mapping at scale

**DOI:** 10.64898/2026.09.20.753039

**Authors:** Hae Been Lee, Junyeong Ma, Han-June Kim, Kyungwoo Song, Insuk Lee

## Abstract

Graph-based protein function analysis is powerful, but protein-protein interaction (PPI) networks exist for only a small fraction of animal and plant genomes. We present LoGoPPI, which infers PPIs from sequence by combining bi-encoder global protein representation with local residue-level late interaction. LoGoPPI matches or exceeds state-of-the-art PLM-based cross-encoders while achieving orders-of-magnitude faster inference, up to ∼1,500-fold, and its local branch provides residue-level signals associated with interaction interfaces and structurally flexible regions. This efficiency enables practical proteome-wide PPI reconstruction at scales prohibitive for cross-encoder models, potentially extending interactome mapping to tens of thousands of animal and plant species. LoGoPPI thus provides a scalable framework for comparative and functional analysis of protein networks across diverse taxa.

## Main

Rapid advances in genome sequencing and assembly are expanding the catalog of proteins across the biosphere, including large and complex animal and plant genomes. The Earth BioGenome Project aims to generate reference genomes for ∼150,000 eukaryotic species by 2030^1^, making comprehensive protein functional annotation a major next challenge. Although homology-based annotations using sequence and structural similarity are highly effective, it remains limited for proteins lacking detectable homologs. Graph-based functional analysis can help address this gap, but experimental PPI mapping at this scale is infeasible, and computationally inferred PPI networks are currently available for only a small fraction of sequenced genomes.

Recent studies have shown that protein language models (PLMs) can infer PPIs directly from sequence and enable reconstruction of genome-scale PPI networks across diverse taxa^2–6^. Existing PLM-based PPI predictors largely follow either a bi-encoder or cross-encoder design. Early methods such as D-SCRIPT^2^, Topsy-Turvy^3^, TT3D^4^, and TUnA^5^ use bi-encoders that encode each protein independently, which are computationally efficient; however, predictive performance remains constrained, and subsequent extensions attempting to incorporate inter-sequence interaction signals have led to only limited gains. In contrast, cross-encoder approaches such as PLM-interact^6^ and PPLM-PPI^7^ jointly encode protein pairs and often achieve higher accuracy, yet their all-to-all token attention is computationally prohibitive at genome scale, restricting practical network reconstruction for most animal and plant species.

To combine the computational efficiency of bi-encoders with the accuracy of cross-encoders, we adopted the late-interaction architecture introduced in ColBERT^8^, which independently encodes the two inputs and then applies a lightweight yet expressive token-level interaction to capture fine-grained similarity. Building on this idea, we propose LoGoPPI, a PPI inference model that integrates bi-encoder “global” embedding similarity with “local” late interaction over residue-level representations of two proteins (**Fig. 1a**).

**Fig. 1.**
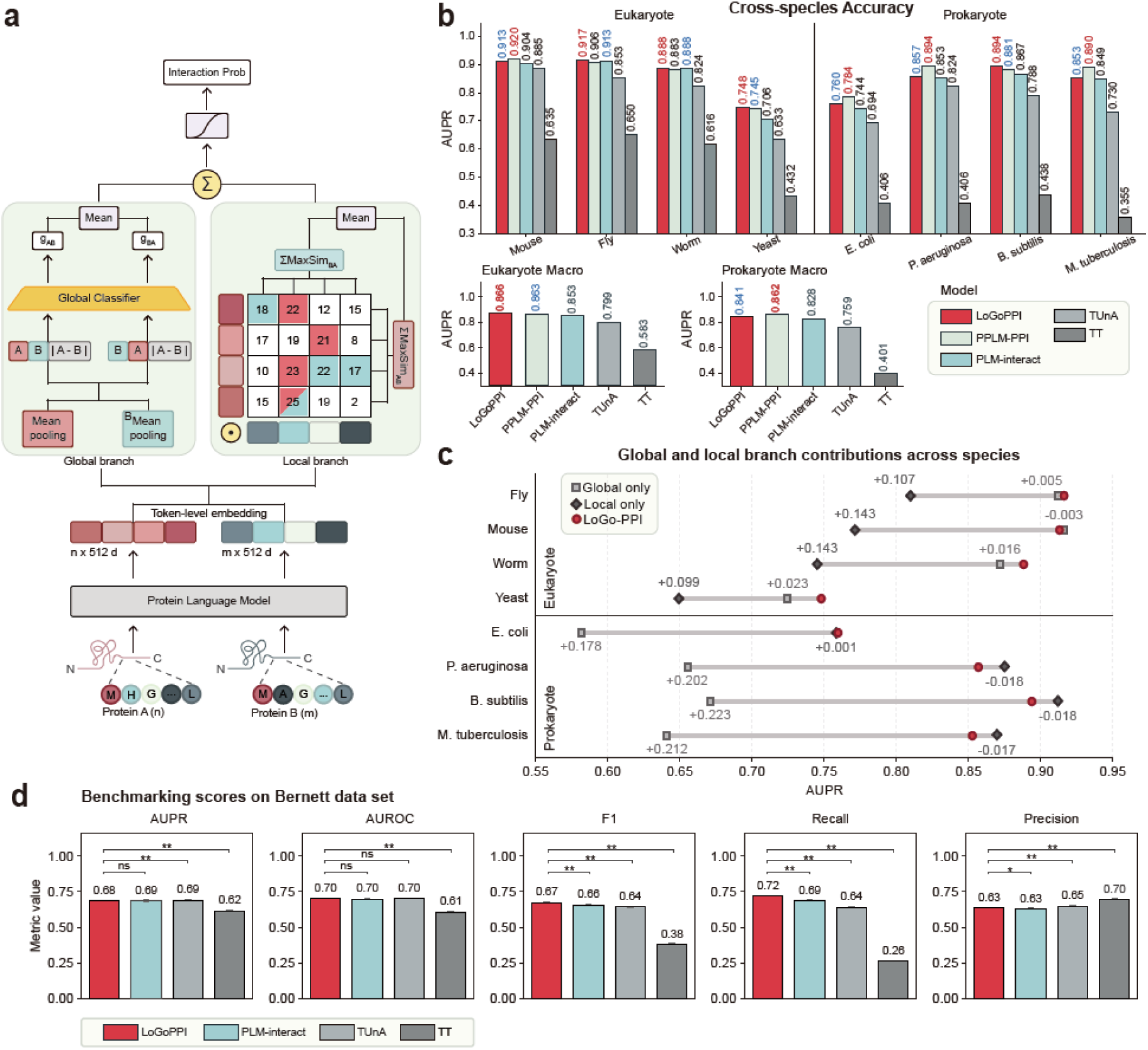
LoGoPPI architecture and benchmark performance. a, Overview of the LoGoPPI architecture. **b,** Cross-species area under the precision–recall curve (AUPR) and macro-AUPR across taxa. **c,** Branch ablation analysis across species. The AUPR changes obtained by adding Local-only branch to the Global-only model and by adding the Global-only branch to the Local-only model are displayed alongside the corresponding markers. **d,** Performance of LoGoPPI and other PLM-based PPI inference models on the leakage-reduced Bernett benchmark.

LoGoPPI processes ESM-2^9^ residue embeddings through complementary global and local branches. The global branch mean-pools residue embeddings into protein-level representations and applies a multilayer perceptron classifier to produce a global logit, whereas the local branch uses a ColBERT-style MaxSim operation, retaining for each residue its maximum similarity to any residue in the partner protein and averaging these maxima to obtain a length-normalized directional interaction score. Because the two branch outputs operate on different numerical scales, the global logit is calibrated by temperature scaling, whereas the MaxSim score is standardized using training-set statistics and then calibrated to a local logit. The calibrated logits are equally combined and transformed by a sigmoid function to obtain the final interaction probability. To ensure invariance to protein input order, both branches compute scores in both directions and average them. Consistent with this symmetric design, reversing the input order did not significantly alter predicted interaction probabilities (**Supplementary Fig. 1**).

This late-interaction design has three advantages: (1) MaxSim captures sparse, binding site– driven signals that pooled embeddings can dilute, (2) its permutation-invariant matching enables alignment-free residue-level comparison without assuming fixed positional alignment of binding site, and (3) it replaces computationally expensive attention with simple max-based residue matching, substantially reducing computational cost^8^.

We compared LoGoPPI with four PLM-based PPI predictors across eight cross-species benchmarks comprising four eukaryotic and four prokaryotic species, derived from the STRING PPI database^10, 11^ (**Fig. 1b**). All models were trained exclusively on human PPIs and evaluated at a 1:10 positive-to-negative ratio, with area under the precision–recall curve (AUPR) as the primary metric. LoGoPPI achieved the highest macro-AUPR across the four eukaryotic benchmarks (0.866), narrowly exceeding PPLM-PPI (0.863), and ranked second across the four prokaryotic benchmarks (0.841), behind PPLM-PPI (0.862); PLM-interact generally showed the third-best performance.

To assess the contribution of each LoGoPPI branch, we performed ablation analysis across the eight cross-species benchmarks (**Fig. 1c**), comparing the full model with Global-only and Local-only variants. In eukaryotes, LoGoPPI performed similarly to Global-only and substantially better than Local-only, indicating that global protein-level representations provided the dominant predictive signal. In contrast, in prokaryotes, LoGoPPI substantially outperformed Global-only, whereas Local-only slightly exceeded the full model, indicating a greater contribution from residue-level interaction signals. These results suggest that eukaryotic PPIs may be better captured by similarities in global protein representations, whereas prokaryotic PPIs may be more effectively characterized by localized residue-level interactions. Despite these distinct branch dependencies, the combined global–local architecture maintained competitive performance across both groups.

We further assessed whether cross-species performance was driven by close sequence similarity to proteins in the human training set. Although AUPR gradually decreased with lower sequence identity for all PLM-based PPI predictors, all models remained substantially above the random baseline even at 20-40% bin. LoGoPPI maintained the highest AUPR across the full sequence-identity range except 0–20%, where it ranked second to PPLM-PPI by only 0.008 AUPR, indicating robust cross-species generalization beyond close human homologs (**Supplementary Fig. 2**).

To evaluate performance under stricter control of train–test leakage, we assessed LoGoPPI on the leakage-reduced human benchmark of Bernett et al.^12^ (**Fig. 1d**). PPLM-PPI was excluded because no checkpoint trained on the Bernett training set was publicly available, whereas corresponding checkpoints were available for PLM-interact and TUnA. LoGoPPI achieved AUPR and AUROC broadly comparable to those of the other models. At a 0.5 classification threshold, it showed significantly higher recall but lower precision than all comparators, yet achieved a significantly higher F1 score than all other models despite this precision–recall trade-off.

We next compared sequence-based PPI inference with structure-based prediction using AlphaFold3 (AF3)^13^ by evaluating LoGoPPI and other PLM-based models on multispecies benchmarks derived from experimentally resolved PDB complexes (**Supplementary Fig. 3**). AF3 achieved the highest AUPR (0.956), followed by PPLM-PPI (0.884), LoGoPPI (0.878), TUnA (0.855), and PLM-interact (0.836). Thus, although AF3 provided superior accuracy on resolved complexes, PLM-based sequence models, particularly LoGoPPI and PPLM-PPI, retained strong performance without the computational cost of explicit structure prediction, supporting their practical utility for large-scale PPI screening.

We further evaluated LoGoPPI, PLM-interact, and TUnA on Bernett hard-negative sets in which protein pairs shared either a Pfam accession or a GO Cellular Component annotation (**Supplementary Fig. 4**). All models performed worse than on size-matched random negatives, indicating difficulty in distinguishing true interactions from biologically similar non-interacting pairs. Nevertheless, LoGoPPI remained comparable to or better than the other PLM-based predictors, with AUPRs of ∼0.64 on both hard-negative benchmarks, above the random expectation of 0.5 for balanced classes. These results underscore the sensitivity of PPI performance to negative-set composition while supporting the relative robustness of LoGoPPI under stringent hard-negative evaluation.

While LoGoPPI achieved predictive performance comparable to or better than existing PLM-based PPI predictors, its major advantage was computational scalability. Using an NVIDIA B200 GPU, we benchmarked all models on the same first 14 million unique pairs among 10,000 proteins sampled from the mouse benchmark. LoGoPPI completed inference in 533.87 s (8.90 min), compared with 9.44 h for TUnA, 5.84 days for PLM-interact, and 9.38 days for PPLM-PPI, corresponding to 63.7-, 945.5-, and 1,518-fold longer runtimes, respectively (**Fig. 2a**). To complement this hardware- and implementation-dependent wall-clock benchmark, we estimated theoretical computational cost in FLOPs as a hardware-independent measure of architectural scaling. This analysis indicated an approximately 2,300-fold lower computational cost for LoGoPPI than the evaluated cross-encoder models at this scale (**Supplementary Fig. 5**). This efficiency enabled proteome-scale inference: LoGoPPI screened all pairwise combinations among ∼45,000 quinoa (*Chenopodium quinoa*) proteins within a day, yielding a network enriched for interactions among heat-responsive genes and for functional and co-expression relationships (**Fig. 2b-c**; **Supplementary Fig. 6**). Together, these results demonstrate that LoGoPPI combines competitive accuracy with orders-of-magnitude faster inference, enabling practical proteome-scale PPI mapping.

**Fig. 2.**
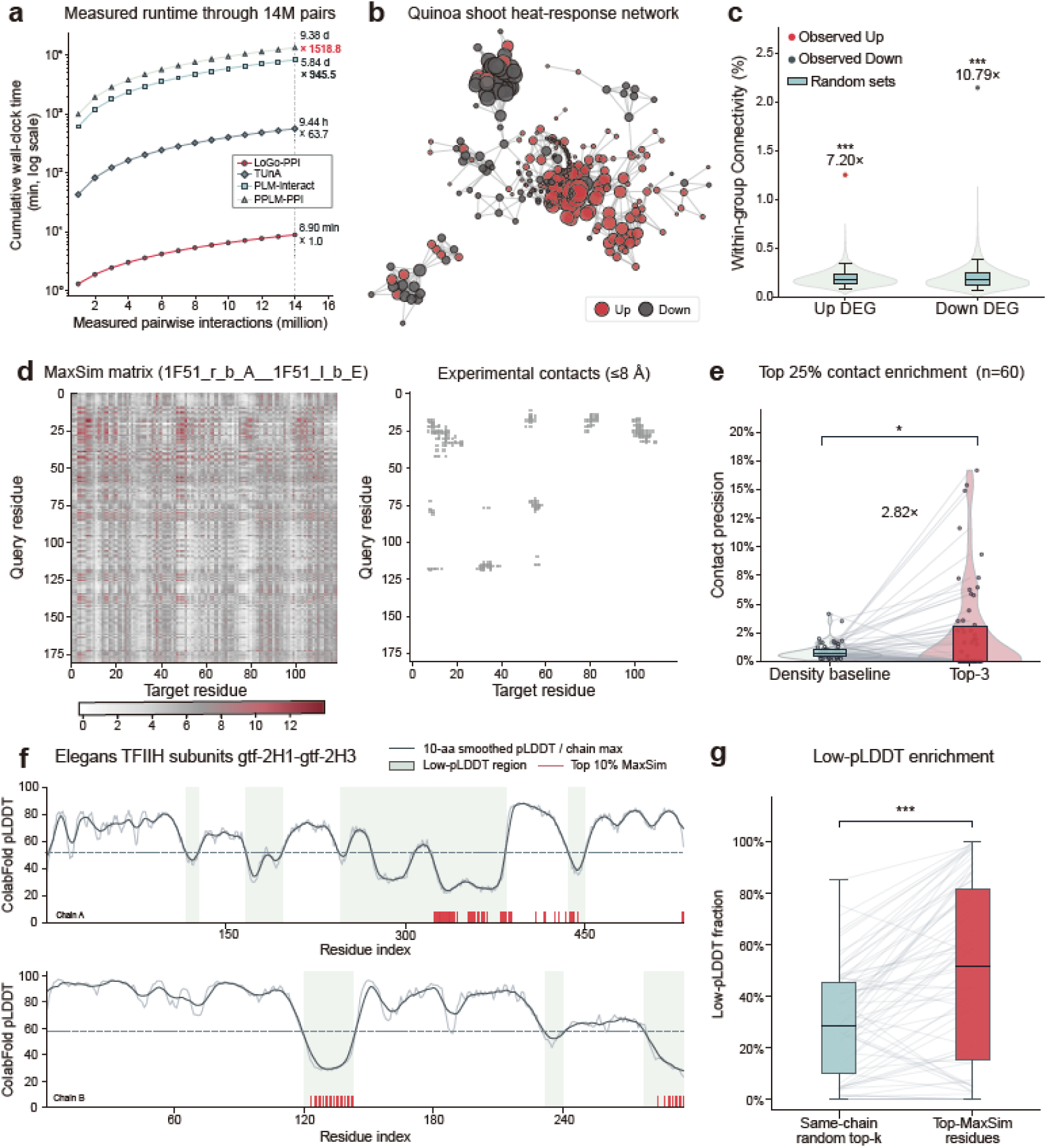
Scalability and interpretability of LoGoPPI. a, End-to-end wall-clock time for 14 million pairs from the cross-species Mouse dataset, with speedup defined relative to LoGoPPI. **b,** Quinoa shoot heat-response network comprising up- and down-regulated quinoa genes derived from quinoa (*Chenopodium quinoa*) interactome. Node size was weighted by the number of interacting proteins. **c,** Within-group connectivity among up- and down-regulated quinoa genes in shoots under heat treatment, compared with a null distribution from 5,000 random gene sets of matched size (one-sided empirical tests, FDR = 0.0002 for both). **d,** MaxSim matrix and experimental contacts (≤8 Å) between the Spo0B phosphotransferase and Spo0F response regulator involved in *Bacillus subtilis* sporulation signaling (PDB: 1F51). **e,** Top-3 MaxSim residue pairs showed higher contact precision than the density-matched baseline across 60 complexes (2.82-fold;*P* = 0.0496). **f,** pLDDT-track analysis: pLDDT profiles of a representative protein pair, showing smoothed pLDDT (dark line), low-pLDDT regions (gray), and Top-MaxSim residues (red). **g,** Top-MaxSim residues were significantly enriched in low-pLDDT regions compared with same-chain random controls across 72 pairs (one-sided paired Wilcoxon test,*P* = 3.51 × 10^−8^). Boxes indicate the interquartile range, and whiskers show the minimum and maximum values.

Beyond computational scalability, LoGoPPI provides residue-level interpretability through its local branch. In experimentally resolved complexes, reciprocal top-MaxSim residue pairs, where each residue ranked the other among its top three matches in the partner protein, frequently overlapped structural interfaces (**Fig. 2d**; **Supplementary Fig. 7**). Among the highest-scoring quartile of complexes, these pairs showed 2.82-fold higher contact precision than expected from the contact-density baseline (*P* = 0.0496; **Fig. 2e**). Top-MaxSim residues were also enriched in low-pLDDT regions, which may reflect structural flexibility, intrinsic disorder, or prediction uncertainty^14, 15^ (**Fig. 2f**; **Supplementary Fig. 8**), occurring in 49.2% of cases versus 29.8% for same-chain random controls (**Fig. 2g**). Although MaxSim is not a direct contact predictor, these findings indicate that it captures local interaction signals associated with protein interfaces and structurally flexible regions.

We also assessed protein domain-associated prediction biases by annotating Bernett test-set proteins with Pfam domains from UniProtKB and analyzing domains represented in at least 50 positive and 50 negative pairs. Domains were classified by their effects on recall and false-positive rate as “improved separation”, “positive-call bias”, “negative-call bias”, or “reversed separation” (**Supplementary Fig. 9**; **Supplementary Table 1**). These results reveal domain-dependent prediction biases in LoGoPPI and can guide interpretation and experimental prioritization of family-specific predictions.

Together, these results establish LoGoPPI as a highly scalable PPI inference framework that combines competitive accuracy with orders-of-magnitude faster computation. Its efficiency enables practical proteome-wide interaction mapping, including in large proteomes, and may support reconstruction of PPI networks across the thousands to tens of thousands of animal and plant species expected from ongoing genome sequencing efforts, facilitating comparative and functional analyses of protein networks across diverse taxa.

## Methods

### Global-local architecture of LoGoPPI

We employed ESM-2^9^ (650M parameters) as the backbone protein language model. ESM-2 was pretrained on large-scale protein sequence data and encodes sequence composition and evolutionary relationships. The pretrained ESM-2 backbone, projection layer, and global classifier were fine-tuned separately for each benchmark dataset for protein-protein interaction (PPI) prediction. The corresponding fine-tuning procedures are described in the “Training details” section.

Given two protein sequences, *A* and *B*, the fine-tuned ESM-2 backbone generates a 1,280-dimensional contextualized representation for each residue. These embeddings are projected to 512 dimensions through a shared linear layer. After excluding BOS, EOS, and padding tokens, the resulting residue-level representations are denoted by

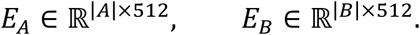

LoGoPPI comprises two complementary scoring branches: a global branch that captures protein-level relationships and a local branch that captures residue-level similarities between two proteins.

For the global branch, mean pooling is applied over the valid residue positions to obtain protein-level representations *p_A_* and *p_B_*.

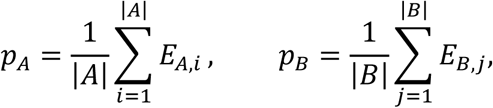

where *p_A_*, *p_B_* ∈ ℝ^512^. For an ordered protein pair (*A*, *B*), the global feature vector is constructed by concatenating the two pooled representations and their element-wise absolute difference:

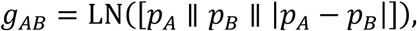

where || denotes vector concatenation and LN denotes layer normalization. The resulting 1,536-dimensional vector is passed through a global classifier implemented as a two-layer multilayer perceptron, consisting of a 1,536-to-512 linear layer, ReLU activation, dropout, and a 512-to-1 output layer, producing the ordered global logit l*_AB_* . Because the concatenated representation depends on the input order, the same classifier is also evaluated for the reversed pair (*B*, *A*). The global branch logit is the average of the two directional logits:

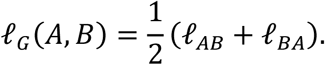

The local branch uses a symmetric MaxSim operation inspired by the late-interaction mechanism of ColBERT^8^. Pairwise dot-product similarities are first computed between all valid residue embeddings in *E_A_* and *E_B_*. For each residue in *A*, the maximum similarity to any residue in *B* is retained and averaged across residues:

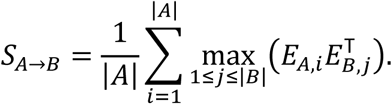

The reverse-direction score *S_B_*_→*A*_ is computed analogously and the symmetric MaxSim score is the average of the two directional scores.

Because the global and local branch outputs operate on different numerical scales, they are calibrated separately. The global logit is transformed using temperature scaling with an additive bias:

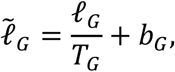

where *T_G_* and *b_G_* are learned calibration parameters estimated on the validation set. Whereas the MaxSim score is first standardized using the training-set mean *μ_train_* and standard deviation *σ_train_*, and then converted to a calibrated local logit:

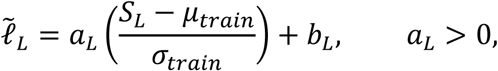

where *a_L_* and *b_L_* are a positive scaling parameter and an additive bias, respectively, fitted on the validation set. The final interaction logit is obtained by equally combining the calibrated global and local logits:

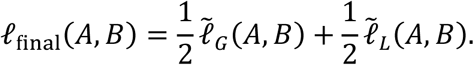

The predicted interaction probability is then given by *y*^ = *σ*(l_final_), where *σ* denotes the sigmoid function.

### Construction of cross-species benchmark datasets

To evaluate cross-species generalization, we used a benchmark dataset based on STRING (v11)^10^, originally introduced by D-SCRIPT^2^. Because this benchmark has been widely used to train and evaluate subsequent PLM-based PPI prediction models, we adopted it to facilitate direct comparison with prior methods. Human protein–protein interactions (PPIs) were used for model training, while test sets were constructed from five additional species: *Mus musculus* (Mouse), *Drosophila melanogaster* (Fly), *Caenorhabditis elegans* (Worm), *Saccharomyces cerevisiae* (Yeast)*, and Escherichia coli.* All datasets contained only high-confidence physical PPIs supported by experimental evidence scores.

Protein sequences were restricted to 50–800 amino acids. To minimize the influence of sequence similarity, redundancy was reduced using CD-HIT^16^ with a 40% sequence identity threshold. Negative pairs were then generated by random pairing at a 10:1 negative-to-positive ratio, reflecting the sparsity of true PPIs in biological systems. For the human dataset, this procedure yielded 47,932 positive PPIs, which were randomly split into 38,345 interactions for training (80%) and 9,587 for validation (20%).

For cross-species evaluation, we used the curated PPLM-PPI benchmark^7^. The original dataset contained 5,000 positive and 50,000 randomly generated negative pairs for each eukaryotic species, and 2,000 positive and 20,000 negative pairs for *E. coli*. Following PPLM-PPI curation, duplicate pairs and pairs with conflicting labels were removed without replacement. The final negative-set sizes were 49,999 for Mouse, 49,993 for Fly, 49,996 for Worm, 49,963 for Yeast, and 16,232 for *E. coli*, while the positive-set sizes remained unchanged.

LoGoPPI used a maximum sequence length of 800 residues for both fine-tuning and inference. Because 800 residues fall within the original sequence-length capacity of ESM-2^9^, we retained the original ESM-2 architecture and rotary positional encoding without modification.

### Construction of additional prokaryotic cross-species benchmarks

To further evaluate cross-species generalization in prokaryotes, we constructed test sets for *Pseudomonas aeruginosa* PAO1 (NCBI taxonomy ID: 208964), *Bacillus subtilis* subsp. *subtilis* strain 168 (224308), and *Mycobacterium tuberculosis* H37Rv (83332) using STRING (v12.0)^11^. Because the experimental evidence channel alone yielded limited coverage, positive pairs were selected from the STRING physical interaction network using a stringent combined confidence score ≥900.

Proteins of 50–1,000 amino acids were retained and clustered within each species using MMseqs2 at 40% sequence identity, with one representative per cluster. Putative negatives were generated by random within-species pairing after excluding known positives, self-pairs, and duplicate undirected pairs, and were sampled without replacement at a 10:1 negative-to-positive ratio. The final datasets contained 1,840 positive and 18,400 negative pairs for *P. aeruginosa*, 1,702 and 17,020 for *B. subtilis*, and 1,525 and 15,250 for *M. tuberculosis*. During LoGoPPI inference, sequences longer than 800 residues were truncated to the first 800 amino acids.

### Evaluation on the leakage-reduced Bernett benchmark

The Bernett benchmark^12^ is a human PPI gold-standard benchmark designed to minimize train-test leakage. Positive PPIs were derived from HIPPIE (v2.3)^17^ and subsequently partitioned using KaHIP, a balanced graph-partitioning algorithm, to reduce topological overlap across training, validation, and test sets. CD-HIT clustering at ≤40% sequence identity was further applied to limit sequence-level leakage between splits. Negative interactions were generated by random pairing of protein pairs not reported in HIPPIE and sampled at a 1:1 ratio relative to positive interactions. A total of 163,192 protein pairs were used for training, 59,260 for validation, and 52,048 for testing. During training, the maximum sequence length was set to 800 residues, and sequences exceeding this length were truncated at the C-terminus.

Model performance was evaluated on the held-out Bernett test set by benchmarking LoGoPPI, PLM-interact, and TUnA under identical conditions; PPLM-PPI was excluded because a checkpoint trained on the Bernett benchmark is not publicly available. To quantify uncertainty in performance estimates and assess differences between models, we performed pair-level bootstrap analysis using 2,000 replicates generated by sampling test pairs with replacement. The same resampled pair indices were applied to all models, enabling paired comparisons. For each model, 95% confidence intervals were calculated from the 2.5th and 97.5th percentiles of the bootstrap distribution of each evaluation metric. Two-sided paired bootstrap *P* values for comparisons between LoGoPPI and each competing model were calculated from the corresponding bootstrap distributions of metric differences. Multiple comparisons were adjusted using the Holm procedure separately within each evaluation metric and threshold policy.

### Training details

LoGoPPI is provided in two dataset-specific variants, LoGoPPI-Cross-species and LoGoPPI-Bernett. Both use the same model architecture, and their corresponding checkpoints are publicly available on Hugging Face.

LoGoPPI-Cross-species was trained using three NVIDIA B200 GPUs. The pretrained ESM-2 backbone, projection layer, and global classification branch were jointly fine-tuned using the AdamW optimizer with binary cross-entropy loss. To account for class imbalance in the human training dataset, a positive-class weight of 10 was applied. The maximum sequence length was set to 800 residues per protein. A per-device batch size of 16 with two gradient accumulation steps was used, resulting in an effective batch size of 96 across the three GPUs. Training used a linear learning-rate scheduler with a warmup ratio of 0.1, a weight decay of 0.01, and a maximum gradient norm of 1.0.

The retained human validation pairs were partitioned into model-selection, branch-calibration, and final-score validation subsets, comprising 50%, 25%, and 25% of the retained pairs, respectively. Training was performed for up to 20 epochs, with early stopping (patience = 3) based on AUPR on the model-selection subset. The checkpoints with the highest AUPR on this subset were selected.

After checkpoint selection, all encoder, projection, and classification-head parameters were fixed. Only the branch-specific calibration parameters were subsequently estimated on the branch-calibration subset. The global and MaxSim calibrators were fitted separately by minimizing binary cross-entropy without class weighting. The resulting calibrated branch logits were combined with equal weights, and the final-score validation subset was used to evaluate the combined score.

LoGoPPI-Bernett was trained using the same architecture and training configuration, except that the positive-class weight was set to 1 to reflect the balanced class distribution of the Bernett dataset. For this variant, the validation set was divided equally into model-selection and calibration subsets. The former was used for early stopping and checkpoint selection, whereas the latter was used to fit the global and MaxSim calibrators following the same procedure described above.

All experiments were conducted using a fixed random seed of 42. The learning rate and a complete list of hyperparameters for both model variants are provided in **Supplementary Table 2**.

### Branch ablation analysis

To evaluate the contributions of the global and local branches, we performed a score-level ablation analysis across the eight cross-species benchmark datasets. We compared three variants: (i) Global-only, using the calibrated global logit; (ii) Local-only, using the calibrated MaxSim logit; and (iii) the full LoGoPPI model, which combines the two calibrated branch logits with equal weights. AUPR was calculated separately for each branch-specific score and for the final combined LoGoPPI score.

### Sequence identity–stratified evaluation

To assess the effect of sequence homology to proteins in the human training set, we stratified test pairs by sequence identity and evaluated LoGoPPI, PPLM-PPI, PLM-interact, and TUnA. For each test protein, MMseqs2 was used to identify the highest sequence identity to any protein in the human training set. The homology level of each test pair was defined as the higher of the two protein-level maximum identities. Test pairs were then grouped into five sequence-identity bins: 0–20%, 20–40%, 40–60%, 60–80%, and 80–100%. Within each bin, AUPR was calculated for each model using only protein pairs scored by all four models. Analyses were performed separately for *E. coli*, Fly, Mouse, Worm, and Yeast, and repeated after pooling test pairs across all five species.

### Structure-based PPI benchmark using AlphaFold 3

We compared LoGoPPI with AlphaFold 3 (AF3)^13^ using the interface predicted Template Modeling (ipTM) score, a widely used measure of predicted interface confidence^18^. Because ipTM reflects confidence in a structural interface rather than an interaction probability, it was treated as a proxy for structure-based PPI confidence.

To reduce potential overlap with AF3 training data, we constructed a multispecies benchmark from experimentally determined PDB complexes released on or after 1 October 2021, after the reported AF3 structural training cutoff of 30 September 2021. Inter-chain contacts were defined as non-hydrogen atoms from different chains within 5 Å in biological-assembly coordinates. We retained non-covalent interactions between distinct proteins from the same species, restricted both proteins to ≤800 amino acids, and excluded ambiguous or engineered structures as well as pairs overlapping publicly available training or validation datasets of the sequence-based predictors.

The final benchmark comprised 100 structurally supported positive pairs and 100 matched negative controls drawn from diverse species. Positive selection enforced protein non-redundancy, with at most one pair per PDB entry or protein-sharing complex group and each protein appearing in only one positive pair. The 100 positives included 19 human, 56 non-human eukaryotic, and 25 prokaryotic pairs. Negative controls were generated by within-species rewiring of the same protein endpoints while excluding known interactions, such that each protein appeared once in the positive set and once in the negative set. Species assignments, selected pairs, and model scores are provided in **Supplementary Table 3**.

AF3, LoGoPPI, PPLM-PPI, PLM-interact, and TUnA were evaluated on the same 200 pairs. AF3 predictions were generated using AlphaFold Server with server-generated MSAs, templates disabled, and a fixed random seed of 42 for all protein pairs. Among the five diffusion samples generated for each pair, the sample with the highest internal ranking score was selected, and its A–B chain_pair_iptm value was used as the AF3 score. Sequence-based models were evaluated using their archived checkpoints and standard inference settings.

AUROC and AUPR were calculated at the pair level. To account for dependencies introduced by shared proteins and matched positive–negative construction, related pairs were grouped into 44 resampling blocks. These blocks were sampled with replacement 10,000 times, with identical bootstrap resamples applied across models to derive 95% confidence intervals and paired comparisons.

### Evaluation on biologically similar hard negatives

We evaluated whether LoGoPPI and the comparator models could distinguish true interactions from challenging Bernett negative pairs sharing either a Gene Ontology Cellular Component (CC) annotation or a Pfam domain.

Among the 52,048 Bernett test pairs (26,024 positives and 26,024 negatives), CC-shared pairs were defined as those in which both proteins shared at least one direct CC annotation. Human GO annotations were obtained from the Gene Ontology Consortium release of 21 May 2026, and GO identifiers and relationships were validated against the 26 July 2026 release of go-basic.obo. We retained valid, positively asserted direct annotations and excluded the CC root, terms in the gocheck_do_not_annotate subset, and protein-containing complex (GO:0032991) and its descendants, because shared complex membership could itself imply physical association.

Pfam-shared pairs were defined independently using Pfam cross-references from UniProtKB release 2026_02, retrieved through the UniProt REST API. A pair was considered Pfam-shared if both proteins contained at least one identical Pfam accession.

These criteria identified 19,518 positive and 13,714 negative CC-shared pairs, and 1,483 positive and 286 negative Pfam-shared pairs. For each hard-negative benchmark, 2,000 bootstrap replicates were generated. In each replicate, positive pairs were sampled with replacement from the corresponding eligible positive pool to match the number of hard negatives, and hard negatives were resampled independently with replacement. Size-matched random-negative controls were sampled with replacement from the full set of 26,024 Bernett negatives. The same resampled positive set was used for both hard-negative and random-negative comparisons within each replicate, and all models were evaluated on identical protein pairs.

AUPR, AUROC, and F1 at a fixed score threshold of 0.5 were calculated for each replicate. Reported values represent the mean across 2,000 bootstrap replicates, with 95% confidence intervals defined by the 2.5th and 97.5th percentiles of the bootstrap distributions. Benchmarking on the Bernett-derived hard-negative sets included LoGoPPI, PLM-interact, and TUnA; PPLM-PPI was omitted because a Bernett-trained checkpoint was not publicly available.

### Evaluation of runtime and scalability

To compare PPI inference efficiency, we evaluated LoGoPPI, PPLM-PPI, PLM-interact and TUnA on pairwise interaction prediction among 10,000 proteins sampled from the Mouse cross-species dataset. All 49,995,000 unique undirected protein pairs were enumerated in a fixed order, and the same first 14,000,000 pairs were evaluated for all models. All experiments were performed on a single NVIDIA B200 GPU with a maximum sequence length of 800 residues, using the released inference pipeline for each model.

To ensure stable large-scale inference and enable progress tracking, we used a common wrapper that loaded each model only once and sequentially repeated input loading, inference, and output saving for 1-million-pair chunks. This approach accommodated the single-pair inference design of the PPLM-PPI code while reducing the risk of memory errors and result loss during PLM-interact output storage. Identical pair order and chunk boundaries were used for all models, and total runtime was calculated as the sum of the end-to-end wall-clock times across all chunks.

For LoGoPPI and TUnA, protein representations were computed once and reused across all chunks, with representation-generation time included once in the reported runtime. In contrast, the cross-encoder models, PLM-interact and PPLM-PPI, performed joint pair encoding and prediction for every protein pair, consistent with their released inference workflows. Reported runtime therefore represents cumulative end-to-end wall-clock time, including representation generation, data loading, pair scoring, and output writing.

### Theoretical computational cost analysis

To complement the empirical wall-clock runtime benchmark, we analytically estimated the theoretical computational cost of LoGoPPI, PPLM-PPI, PLM-interact, and TUnA in terms of floating-point operations (FLOPs). Whereas wall-clock time depends on hardware, software optimization, batching, and I/O, FLOP analysis provides a hardware-independent measure of the computational scaling inherent to each model architecture.

For a proteome containing *N* proteins, the number of unique undirected protein pairs was defined as *N*(*N*−1)/2. A multiply–add operation was counted as two FLOPs, and the maximum protein sequence length was fixed at 800 residues for all models.

FLOPs were estimated from the model architecture and inference procedures used in the empirical evaluation. Total computational cost was defined as the sum of protein representation generation and pairwise interaction scoring. For LoGoPPI and TUnA, protein representations are precomputed and reused across interaction predictions, so embedding costs were counted once per protein. In contrast, PLM-interact and PPLM-PPI process each protein pair jointly through a protein language model, and therefore incur the corresponding forward-pass cost for every pair. Additional interaction modules, prediction heads, ensemble evaluations, and bidirectional inference steps were included according to each model’s inference implementation.

Transformer FLOPs were estimated from sequence length, hidden dimension, feed-forward dimension, attention operations, and number of layers. Total costs were calculated for complete all-versus-all proteome graphs ranging from 50 to 100,000 proteins, including both protein-level representation costs and pairwise scoring costs at each proteome size.

### Construction of the quinoa interactome

Protein sequences of *Chenopodium quinoa* were obtained from the Phytozome database (JGI, v1.0 annotation)^19^, comprising a total of 44,776 proteins. LoGoPPI scores were computed for all unique undirected protein pairs, excluding self-pairs. Sequences longer than 800 amino acids were truncated before embedding, and pairs with LoGoPPI scores ≥0.9 were retained as predicted interactions. The resulting network contained 634,152 interactions among 25,400 proteins, corresponding to 56.7% of the quinoa proteome and 0.063% of all possible protein pairs (**Supplementary Fig. 10**).

### Functional validation of the quinoa interactome

We evaluated whether the LoGoPPI-predicted quinoa interactome exhibited two properties commonly observed among physically interacting proteins: functional similarity and transcriptomic co-expression^20^.

### (1) GO term-sharing analysis

Direct GO annotations for transcripts from the *Chenopodium quinoa* ASM168347v1 assembly were retrieved from Ensembl Plants release 63 using Ensembl BioMart^21^. Alternative identifiers and obsolete terms with a single unambiguous replacement were mapped to current GO identifiers. GO terms and ontology assignments were validated against the 26 July 2026 release of go-basic.obo^20^. Invalid annotations, terms with inconsistent ontology assignments, ontology roots, and terms designated by the GO Consortium as unsuitable for direct annotation were excluded.

Biological Process (BP), Molecular Function (MF), and Cellular Component (CC) were analyzed separately. Within each ontology, a protein pair was considered evaluable when both proteins had at least one valid annotation and was classified as GO-sharing if the two proteins shared at least one directly assigned GO term.

GO-sharing enrichment was assessed by comparing the observed LoGoPPI network, defined by scores ≥ 0.9, with 100 random networks. Each random network contained 634,152 edges sampled uniformly from all possible undirected pairs among the same protein set. For each ontology, we calculated the proportion of evaluable edges sharing at least one GO term and defined fold enrichment as the observed proportion divided by the mean proportion across the random networks. One-sided empirical *P* values were calculated as 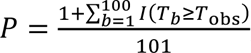, where *T*_obs_ denotes the GO-sharing proportion in the observed network and *T_b_* denotes the corresponding proportion in the *b*th random network, and the three prespecified tests for BP, MF, and CC were adjusted jointly using the Benjamini–Hochberg procedure.

### (2) Co-expression analysis

Twenty-four paired-end RNA-sequencing libraries from quinoa shoots in GSE128155^22^ were obtained from the European Nucleotide Archive. The dataset included four conditions: control, root-only heat treatment, shoot-only heat treatment, and combined root-and-shoot heat treatment, measured at days 1 and 11 with three biological replicates per condition–time combination. RNA-seq read quality was assessed with FastQC (v0.11.7), reads were trimmed using Trimmomatic (v0.38)^23^, and transcript-level TPM values were quantified against the JGI/Phytozome^19^ *C. quinoa* reference transcriptome using Kallisto (v0.44.0)^24^. TPM values from all 24 libraries were combined into a transcript-level expression matrix without gene-level aggregation.

Transcripts were retained if they had TPM ≥1 in at least three samples and nonzero variance after transformation. The three biological replicates within each condition–time combination were then averaged, yielding an eight-dimensional expression profile per transcript. Pairwise co-expression was quantified using the Pearson correlation coefficient across these eight profiles. An interaction was considered evaluable when both transcripts passed the expression filter and was classified as co-expressed when its Pearson correlation was *r* ≥ 0.7.

Co-expression enrichment was evaluated against the same 100 random networks used for the GO analysis. Fold enrichment was defined as the observed proportion of co-expressed evaluable edges divided by the mean proportion across the random networks, with one-sided empirical *P* values calculated as described above, with *T*_obs_ and *T_b_* representing the proportions of co-expressed interactions in the observed network and the *b*th random network, respectively. Co-expression was treated as a separate prespecified test.

To examine whether co-expression increased with LoGoPPI score, all 1,002,422,700 candidate protein pairs were divided into fixed-width score bins of 0.1. The complete score set was scanned sequentially, retaining only pairs for which both transcripts passed the expression filter, and up to 100,000 pairs were sampled without replacement from each bin.

For each score bin, the median Pearson correlation coefficient was used as the primary co-expression measure. Its 95% confidence interval was estimated from 1,000 pair-level bootstrap resamples. To estimate the random expectation within each score bin, the first protein in each sampled pair was retained while the second proteins were randomly permuted among pairs; self-pairs were excluded. This randomization was repeated 100 times per score bin.

### Heat-responsive DEG subnetwork analysis

Heat-responsive genes were identified from the shoot heat-treatment samples in GSE128155^22^ at day 1. Differentially expressed genes (DEGs) were identified using Sleuth based on the estimated effect size and multiple-testing-adjusted significance with |*b*| > 2 and *q* < 0.05. A DEG-specific subnetwork was then extracted from the high-confidence quinoa interactome by retaining only interactions between DEG pairs. Isolated DEGs were excluded from visualization, and subnetworks were displayed using a Kamada–Kawai layout, with nodes colored by the direction of expression change and scaled by degree within the DEG subnetwork.

Connectivity was evaluated separately for upregulated and downregulated DEGs as the proportion of observed interactions among all possible within-group gene pairs. Statistical significance was assessed against null distributions generated from 10,000 randomly sampled gene sets matched in size to each DEG group. Empirical *P* values were adjusted using the Benjamini–Hochberg procedure.

### Structural evaluation of MaxSim residue-pair similarities

To investigate whether residue-level similarities captured by MaxSim corresponded to protein– protein interaction interfaces, we compared MaxSim residue-pair scores with structural contact maps derived from bound complexes in Protein–Protein Docking Benchmark 5.5 (DB5.5)^25^.

MaxSim matrices were indexed according to the model-input sequences, whereas structural residues were indexed according to residues with resolved Cα coordinates in the corresponding PDB structures. Of 271 protein pairs initially considered, 22 were excluded owing to a mismatch between MaxSim matrix dimensions and the number of structurally resolved receptor and ligand residues, and nine further pairs lacking any inter-chain Cα–Cα contact within 8 Å were excluded, leaving 240 structurally evaluable complexes. These complexes were scored using LoGoPPI and ranked by interaction probability; the primary analysis was restricted to the highest-scoring quartile (60 pairs) to test whether MaxSim’s structural signal held among the interactions LoGoPPI predicted with the greatest confidence.

For each complex comprising proteins *A* and *B*, residue embeddings were generated independently using the trained LoGoPPI encoder and projection layer. A residue–residue MaxSim matrix *S* was constructed as

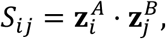

Where 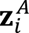 *and* 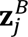 denote the projected embeddings of residues *i* and *j* , respectively. A residue pair (*i*, *j*) was defined as a reciprocal top-three pair when residue *j* was among the three highest-scoring residues in protein *B* for residue *i* in protein *A*, and residue *i* was simultaneously among the three highest-scoring residues in protein *A* for residue *j* in protein *B*.

Two residues from different chains were classified as being in contact when the minimum inter-atomic distance was ≤ 8 Å. Within each complex, reciprocal top-three pairs were ranked by MaxSim score, and the *n*_eval_ = min (*n*_reciprocal_, *n*_contact_) highest-ranking pairs were selected as an interface-size-matched evaluation set, where *n*_reciprocal_ and *n*_contact_ denote the numbers of reciprocal pairs and structural contacts, respectively. The observed contact fraction was *f*_observed_ = *n*_matched_ */ n*_eval_, where *n*_matched_ is the number of evaluated pairs coinciding with structural contacts. The expected contact fraction under uniform random pair selection was *f*_expected_ = *n*_contact_ / (*L_A_* × *L_B_*), with *L_A_* and *L_B_* the numbers of structurally resolved residues in each protein. Observed and expected contact fractions were compared across the 60 complexes using a one-sided paired Wilcoxon signed-rank test.

### Enrichment of high-MaxSim residues in low-pLDDT regions

Protein–protein interactions are frequently mediated by intrinsically disordered regions, short linear motifs, and conditionally structured regions that fold upon binding; such regions often exhibit low pLDDT in single-protein structure predictions^14, 15^. We therefore tested whether residues with high LoGoPPI MaxSim scores were preferentially located in low-pLDDT regions, which would suggest that the model captures sequence features associated with structurally flexible or conditionally structured interaction sites.

Protein pairs were drawn from the cross-species PPI benchmark and restricted to pairs with ≤40% sequence identity to the human training set. Mouse was excluded because too few positive pairs met this criterion. Equal numbers of positive and negative pairs were sampled from *E. coli*, Fly, Worm, and Yeast, yielding 288 protein pairs. The analysis was performed on the top quartile of pairs ranked by LoGoPPI interaction score.

For each residue, the MaxSim score was defined as its maximum embedding similarity to any residue in the partner protein. Within each chain, residues in the top 10% of MaxSim scores were designated Top-MaxSim residues.

Residue-level pLDDT values were obtained from ColabFold-Multimer predictions and smoothed using a centered 10-residue moving average, truncated at chain termini. Smoothed pLDDT values were normalized to the maximum value within each chain, and residues with normalized pLDDT <0.6 were classified as low-pLDDT. Low-pLDDT segments separated by fewer than 30 residues were merged into a single region.

For each chain, enrichment was quantified as the fraction of Top-MaxSim residues located within low-pLDDT regions. A random baseline was generated by sampling the same number of residues without replacement 1,000 times and calculating the corresponding mean fraction. Values for the two chains were averaged within each protein pair. Observed and random fractions were compared across pairs using a one-sided paired Wilcoxon signed-rank test, and fold enrichment was calculated as the ratio of the mean observed fraction to the mean random fraction.

### Protein domain–associated prediction bias analysis

To assess whether LoGoPPI predictions showed systematic bias toward specific protein domains, we evaluated domain-associated differences in recall and false-positive rate (FPR) using the balanced Bernett test set. Protein domains were annotated using Pfam assignments retrieved from UniProtKB. Exact Pfam accessions were retained without hierarchical propagation or merging of related families. Pfam annotations were available for 96.5% of the 3,022 unique proteins in the benchmark, and a protein pair was considered associated with a domain if either protein contained the corresponding Pfam annotation.

The analysis was restricted to domains represented by at least 50 positive and 50 negative protein pairs. LoGoPPI scores were converted to binary predictions using the prespecified probability cutoff of 0.5. For each domain, recall among positive pairs containing that domain was compared with recall among positive pairs without it. FPR was evaluated analogously among negative pairs. Effect sizes were defined as ΔRecall = Recall*_Target_*__*term*_ − ^Recall^*Ot*ℎ*er*_*terms* ^and ΔFPR = FPR^*Target*_*term* ^− FPR^*Ot*ℎ*er*_*terms*.

Confidence intervals were estimated using 2,000 class-stratified bootstrap replicates. Direction-specific differences were tested using one-sided Fisher’s exact tests, with Benjamini–Hochberg correction applied separately to the recall and FPR analyses. Domains with an FDR <0.05 were considered to show a significant association with the corresponding prediction metric.

Protein domains were then categorized according to their joint recall and FPR patterns. A significant increase in recall without a significant increase in FPR was classified as “improved separation.” Significant increases in both recall and FPR indicated “positive-call bias,” whereas significant decreases in both measures indicated “negative-call bias.” A significant decrease in recall together with a significant increase in FPR was classified as “reversed separation.”

For visualization, up to four representative domains were selected from each category. Domains with higher AUROC were prioritized for the better-separation category, whereas those with lower AUROC were prioritized for the poor-or-reversed-separation category. For the positive- and negative-call-bias categories, domains with larger combined absolute effect sizes, defined as |ΔRecall| + |ΔFPR|, were prioritized.

## Code availability

The code in this study is publicly available and has been deposited in GitHub at https://github.com/netbiolab/LoGoPPI. Trained model checkpoints and datasets used in this study are available at Hugging Face. (https://huggingface.co/netbiolab/LoGoPPI-Cross-species; https://huggingface.co/netbiolab/LoGoPPI-Bernett)

## Acknowledgements

This research was supported by the National Research Foundation funded by the Korea Ministry of Science and ICT (RS-2025-18362970, RS-2026-25520168, and RS-2026-25549861 to I.L.). The work was supported in part by Brain Korea 21(BK21) FOUR program.

## Author contributions

H.B.L., and I.L. conceived this study. H.B.L. constructed models and performed evaluations. J.M. and H-J.K. provide advice on data analysis. K.S. provides advice on model design and evaluation. H.B.L. and I.L. wrote the manuscript. All authors read and approved the final manuscript.

## Competing of interests

H-J.K. and I.L. are co-founders of and shareholders in DECODE BIOME Co., Ltd. The other authors declare no competing interests.

## Supplementary Figures

**Supplementary Fig. 1:**
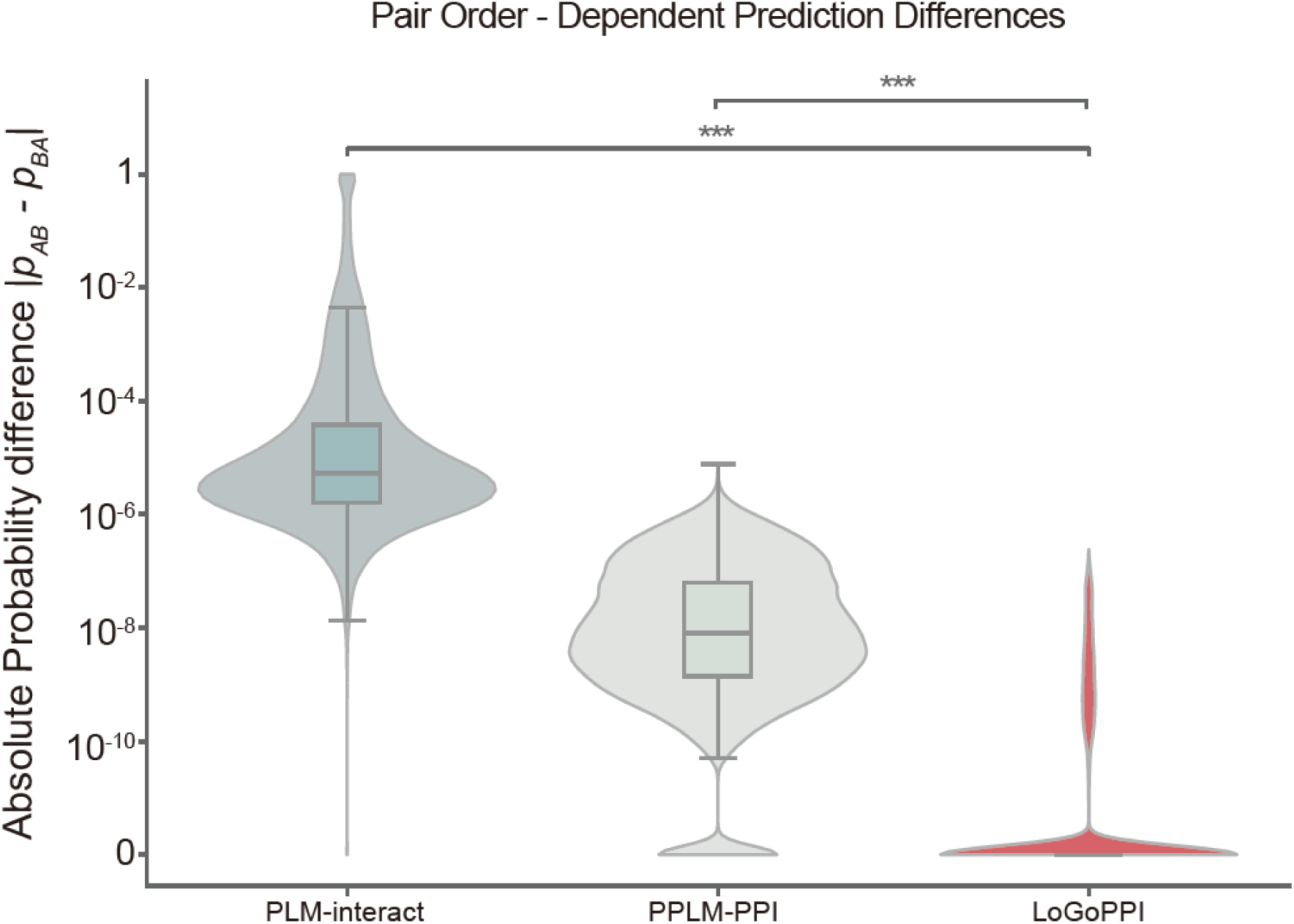
Effect of input pair order on model predictions. Absolute differences in predicted interaction scores between the original and reversed protein-pair orders were evaluated using the Fly dataset from the cross-species benchmark. LoGoPPI showed significantly greater order consistency, reflected by smaller score differences, than PLM-interact and PPLM-PPI (two-sided paired Wilcoxon signed-rank tests with Benjamini– Hochberg correction; both *p*_adj_ < 10^−300^). Violin plots show the full distributions, with embedded box plots indicating the median and interquartile range.

**Supplementary Fig. 2:**
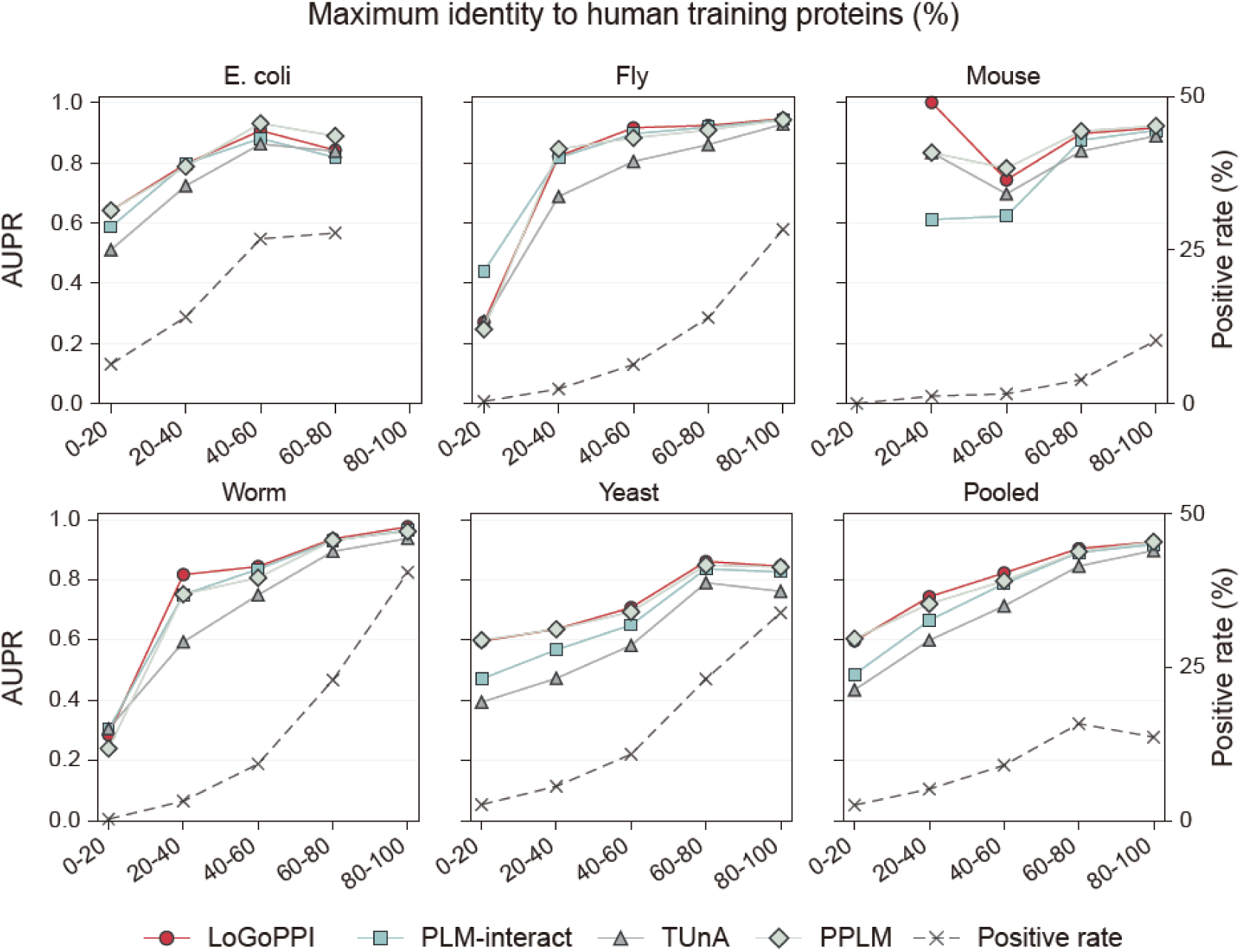
Sequence-homology-stratified cross-species PPI prediction performance. Test protein pairs from *E. coli*, Fly, Mouse, Worm, and Yeast were divided into five bins according to their maximum sequence identity to proteins in the human training set: [0–20], (20–40], (40–60], (60–80], and (80–100] %. For each pair, the maximum identity was defined as the larger of the two protein-level maximum identities. The AUPR was calculated within each bin for LoGoPPI, PLM-interact, TUnA, and PPLM-PPI using only the ordered protein pairs scored by all four models. Species-specific results and results pooled across all five species are shown. Solid lines report AUPR on the left axis, and the dashed line reports the percentage of positive pairs in each bin on the right axis.

**Supplementary Fig. 3:**
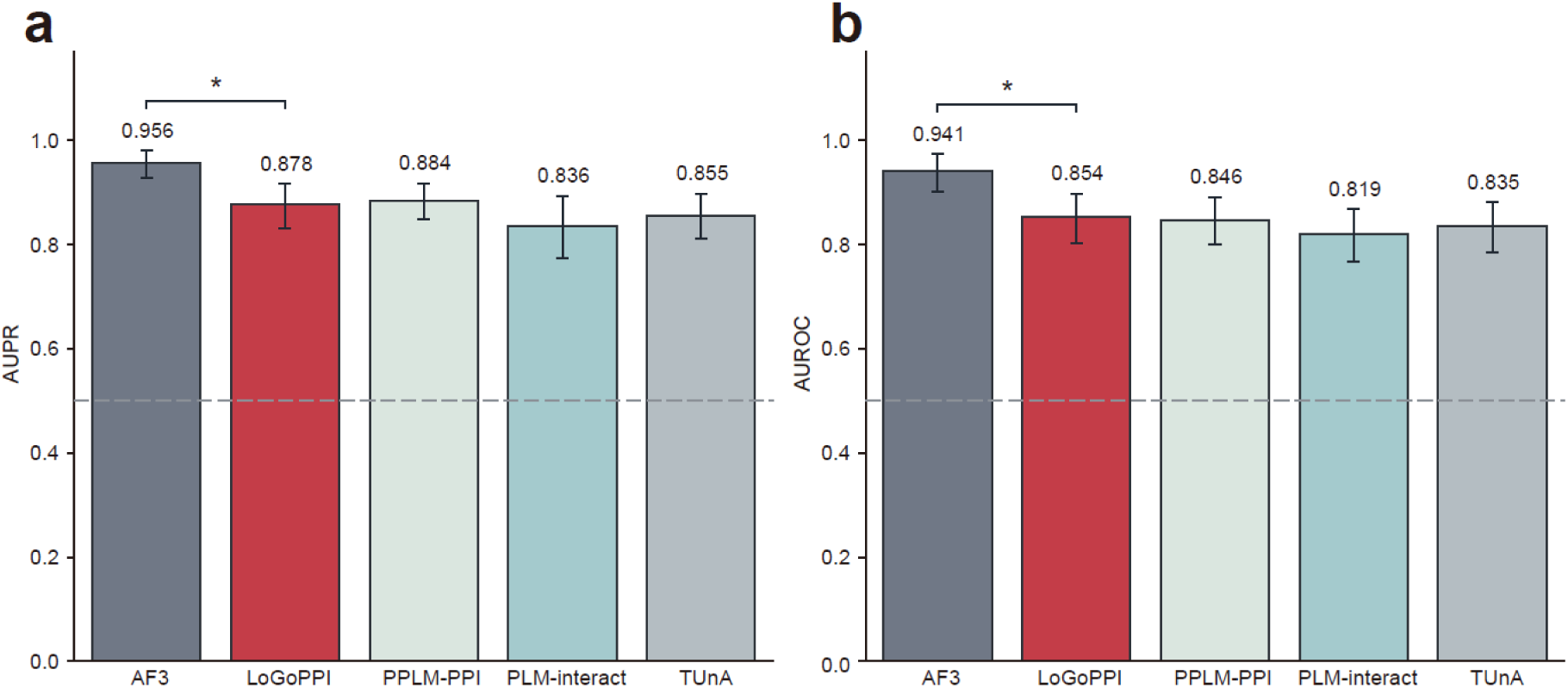
Comparison of Structure- and sequence-based PPI predictions on a multispecies PDB benchmark. The benchmark contains 100 PDB-supported positive pairs and 100 matched presumed-negative pairs from 21 species. **a,** AUPR (average precision). **b,** AUROC. Bars show performance estimates; error bars indicate 95% confidence intervals. Brackets compare AF3 with LoGoPPI (*, the 95% confidence interval for the paired difference excludes zero). Dashed lines mark the 0.5 reference level.

**Supplementary Fig. 4:**
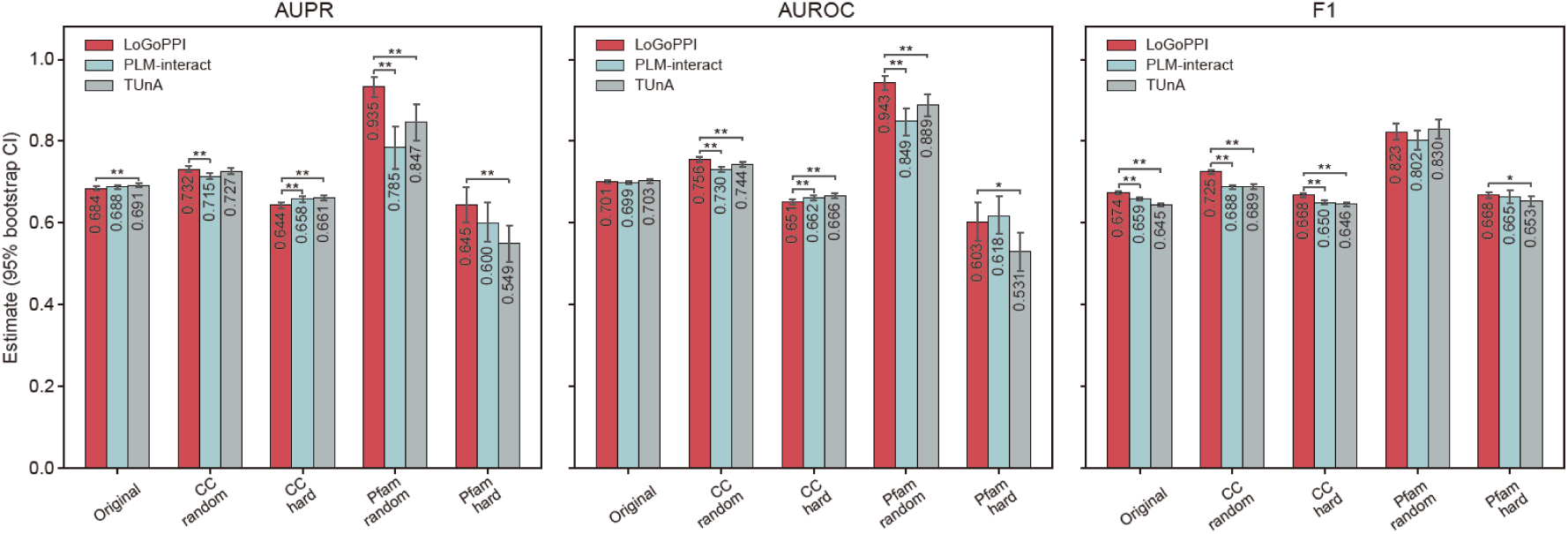
Evaluation on GO- and Pfam-based hard negatives. AUPR, AUROC, and F1 scores are shown for LoGoPPI, PLM-interact, and TUnA on the original test set and on size-matched random-negative and hard-negative subsets constructed according to shared cellular-component (CC) or Pfam annotations. Error bars indicate 95% confidence intervals obtained from 2,000 paired bootstrap replicates. Brackets indicate two-sided paired-bootstrap comparisons between LoGoPPI and each external model. P-values were adjusted across the 30 displayed comparisons using the Benjamini–Hochberg false-discovery-rate procedure. Adjusted significance levels are denoted by *q* < 0.05 (\**), q < 0.01 (**)*.

**Supplementary Fig. 5:**
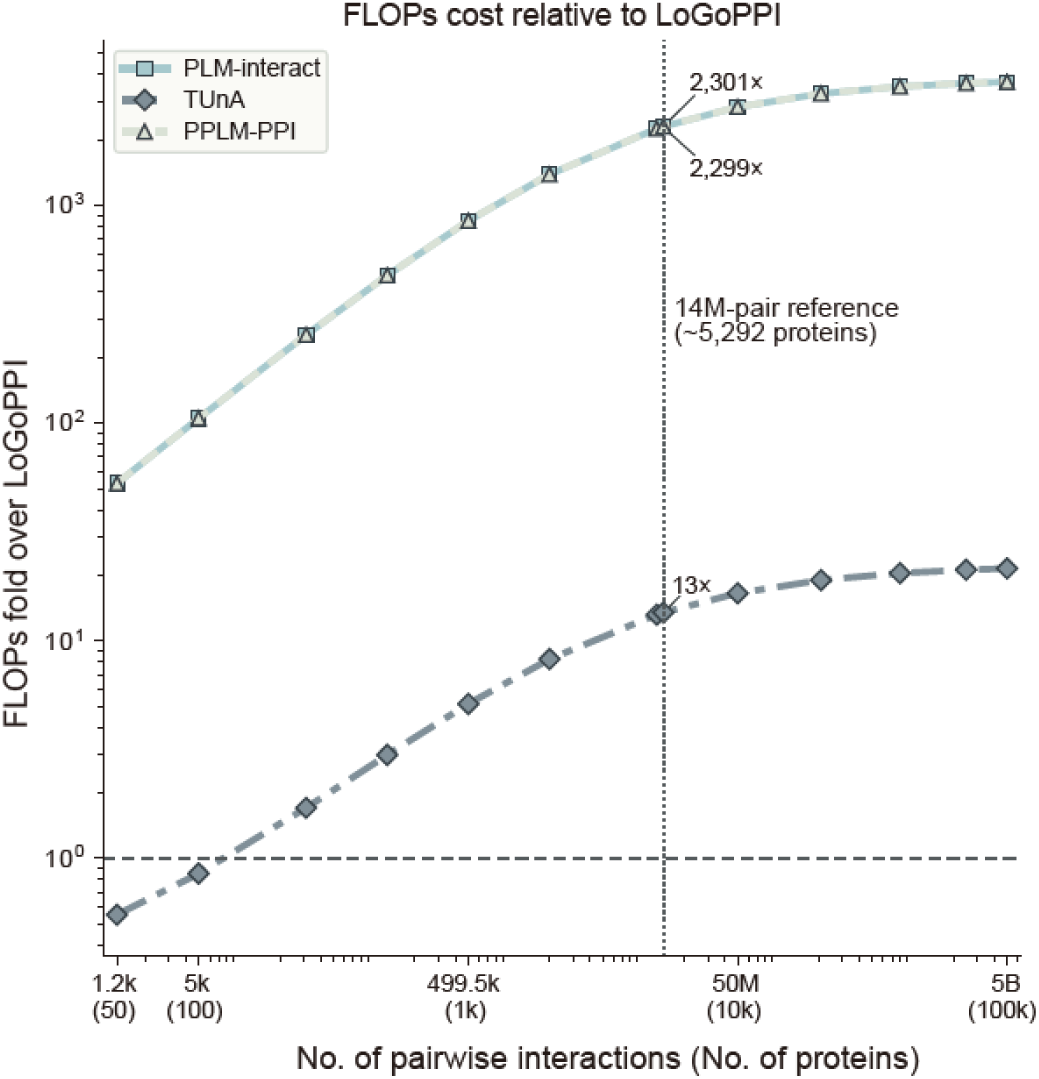
FLOP-based computational complexity analysis. Estimated FLOPs for PLM-interact, TUnA, and PPLM-PPI are shown as fold changes relative to LoGoPPI across increasing numbers of protein pairs and proteins. Both axes are log-scaled, and the horizontal dashed line indicates equal computational cost to LoGoPPI. The vertical dotted line marks the ∼14-million-pair scale used in the empirical runtime benchmark.

**Supplementary Fig. 6:**
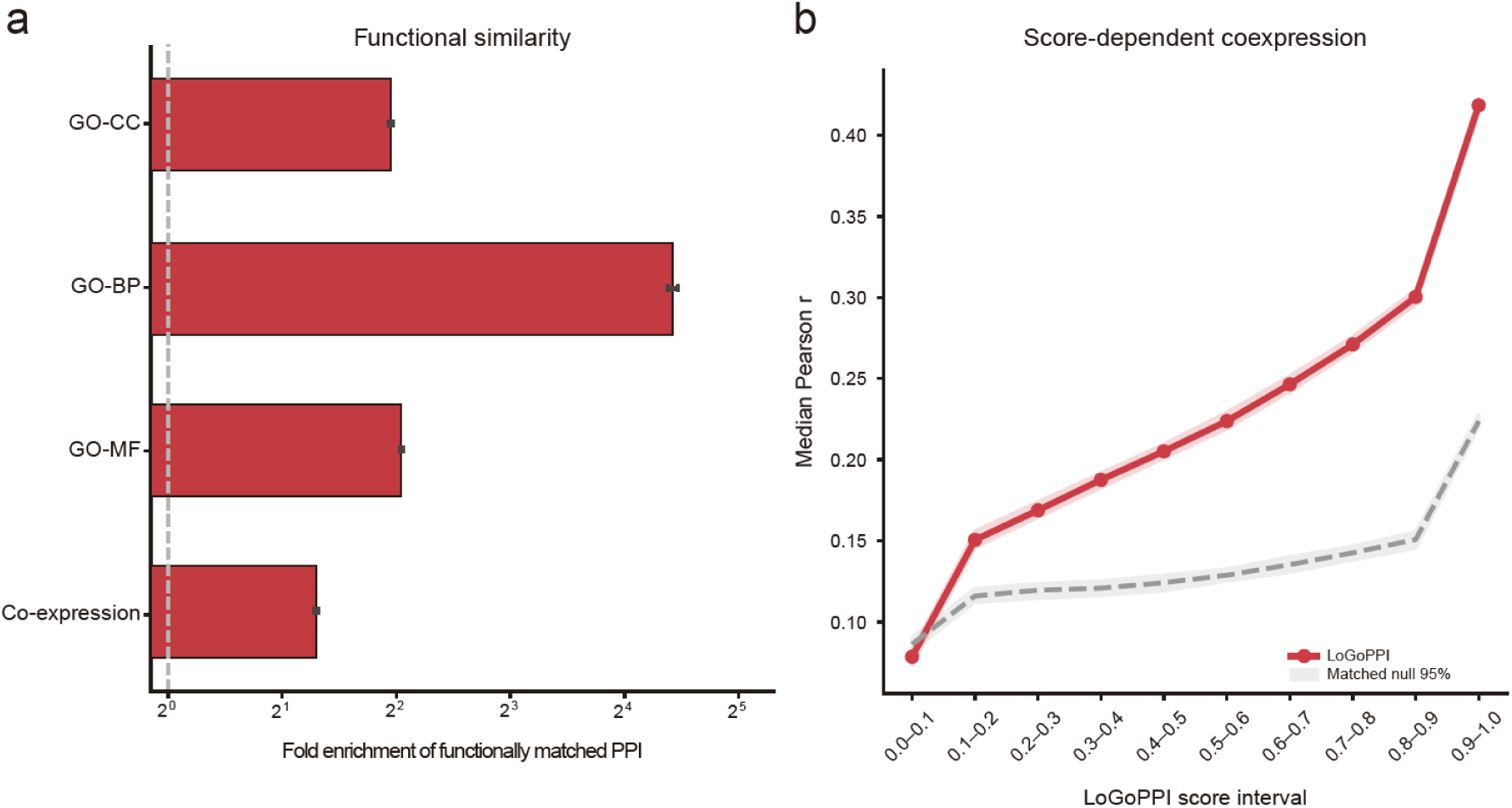
Functional validation of the predicted quinoa PPI network. a, Enrichment of functional concordance in the LoGoPPI network relative to the null model. Bars show observed-to-null ratios for shared cellular compartment, biological process, molecular function, and co-expression; error bars indicate 95% null intervals, and the dashed vertical line marks no enrichment. **b,** Median gene-expression correlation across LoGoPPI score interval. The red line and shading represent the observed network and its 95% confidence intervals, whereas the gray dashed line and band represent the matched-null expectation. Co-expression increased significantly with prediction score, supporting greater functional coherence among higher-scoring protein pairs.

**Supplementary Fig. 7:**
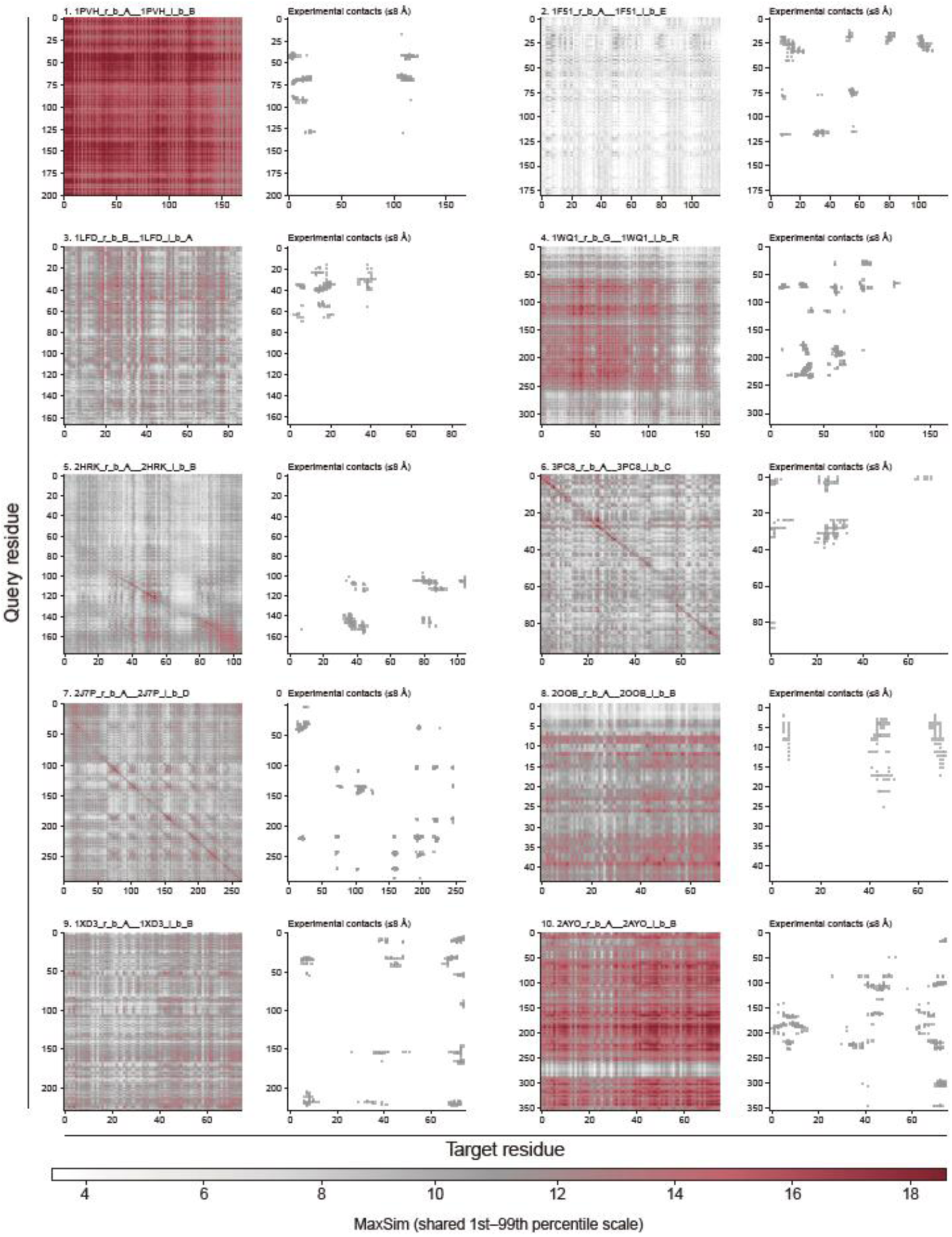
Residue-level MaxSim patterns and experimental contacts for representative protein pairs. Ten protein pairs are ranked by reciprocal Top-3 contact precision. For each pair, the MaxSim residue-similarity matrix is shown alongside the corresponding experimental inter-chain contact map, with contacts defined at a distance of ≤8 Å. Darker red regions indicate higher MaxSim values on a shared color scale. Pair labels report the final LoGoPPI score, observed contact precision, and the number of true-positive residue matches among evaluated matches. The correspondence between high-similarity residues and experimental contacts provides evidence of sparse interaction-site localization and should not be interpreted as dense contact-map prediction.

**Supplementary Fig. 8:**
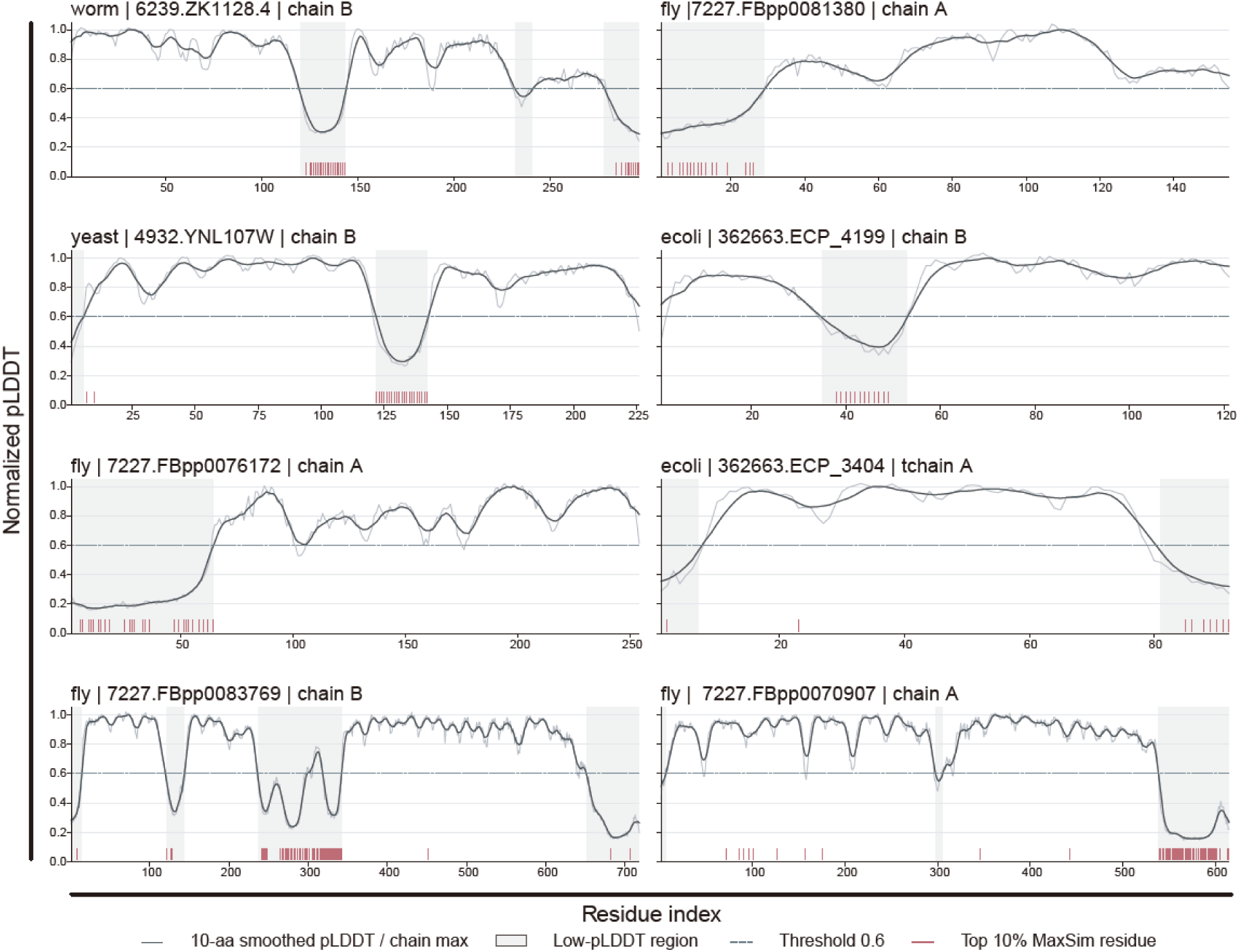
Representative examples of the association between residue-level MaxSim signals and structural confidence. Normalized ColabFold-Multimer pLDDT profiles along the sequence are shown for the eight protein chains with the largest enrichment differences between Top-MaxSim residues and the remaining residues. The dark line indicates the smoothed pLDDT profile; the dashed line marks the low-confidence threshold (normalized pLDDT = 0.6); gray shading highlights low-pLDDT regions. Red ticks indicate Top-MaxSim residues (top 10% of MaxSim scores within each chain).

**Supplementary Fig. 9:**
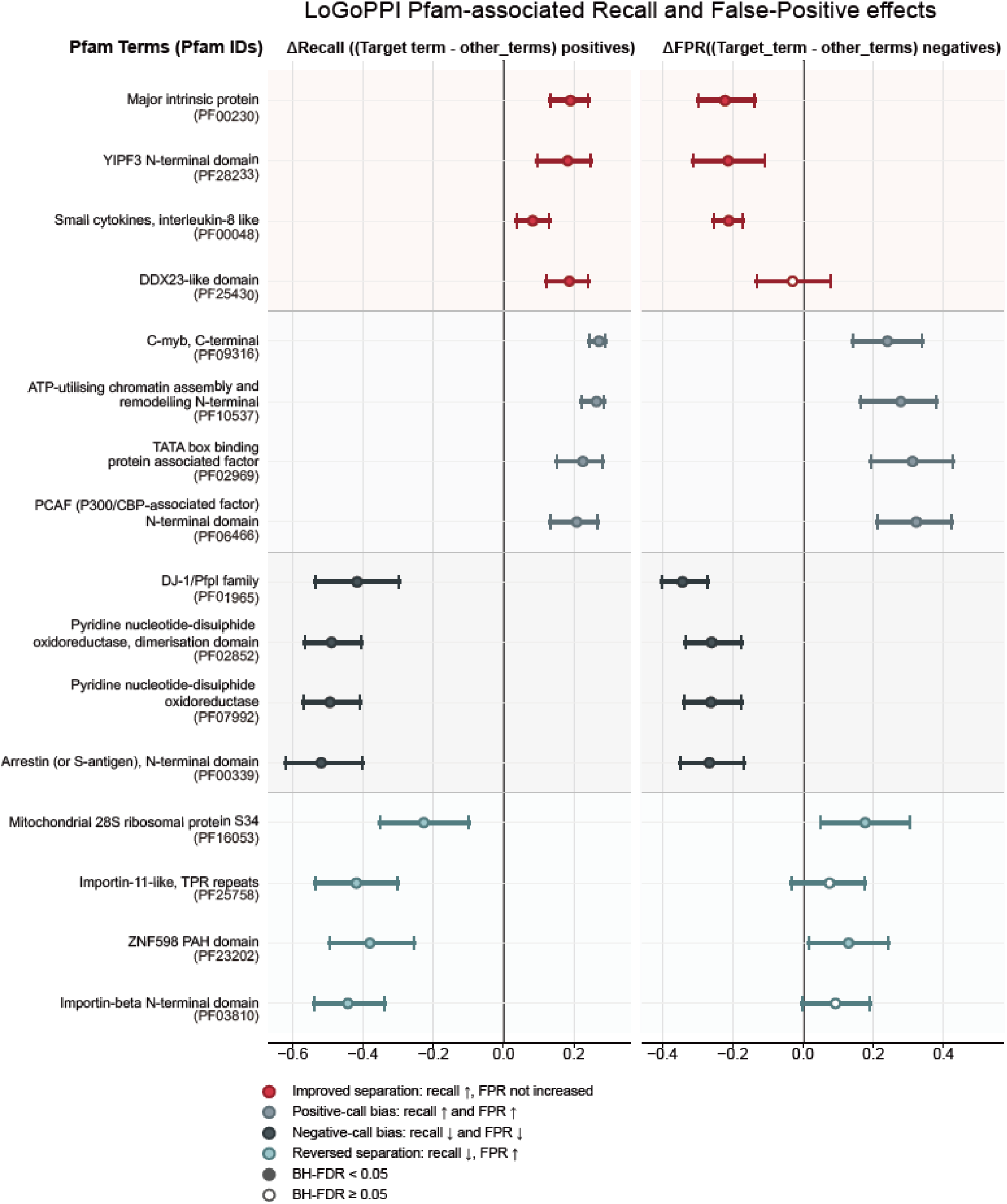
Pfam-associated recall and false-positive effects of LoGoPPI on the Bernett benchmark. Differences in recall between positive protein pairs containing each Pfam and those without that Pfam. Corresponding differences in false-positive rate (FPR) among negative pairs. Positive values indicate higher recall or FPR in Pfam-containing pairs, whereas negative values indicate lower values. Points show estimated differences and horizontal lines indicate 95% confidence intervals. Colors distinguish patterns of improved separation, positive-call bias, negative-call bias, and poor or reversed separation. Filled points indicate recall or FPR differences that were significant in the corresponding direction based on one-sided Fisher’s exact tests with Benjamini–Hochberg false-discovery-rate correction across all eligible Pfams (*q* < 0.05); open points indicate *q* ≥ 0.05.

**Supplementary Fig. 10:**
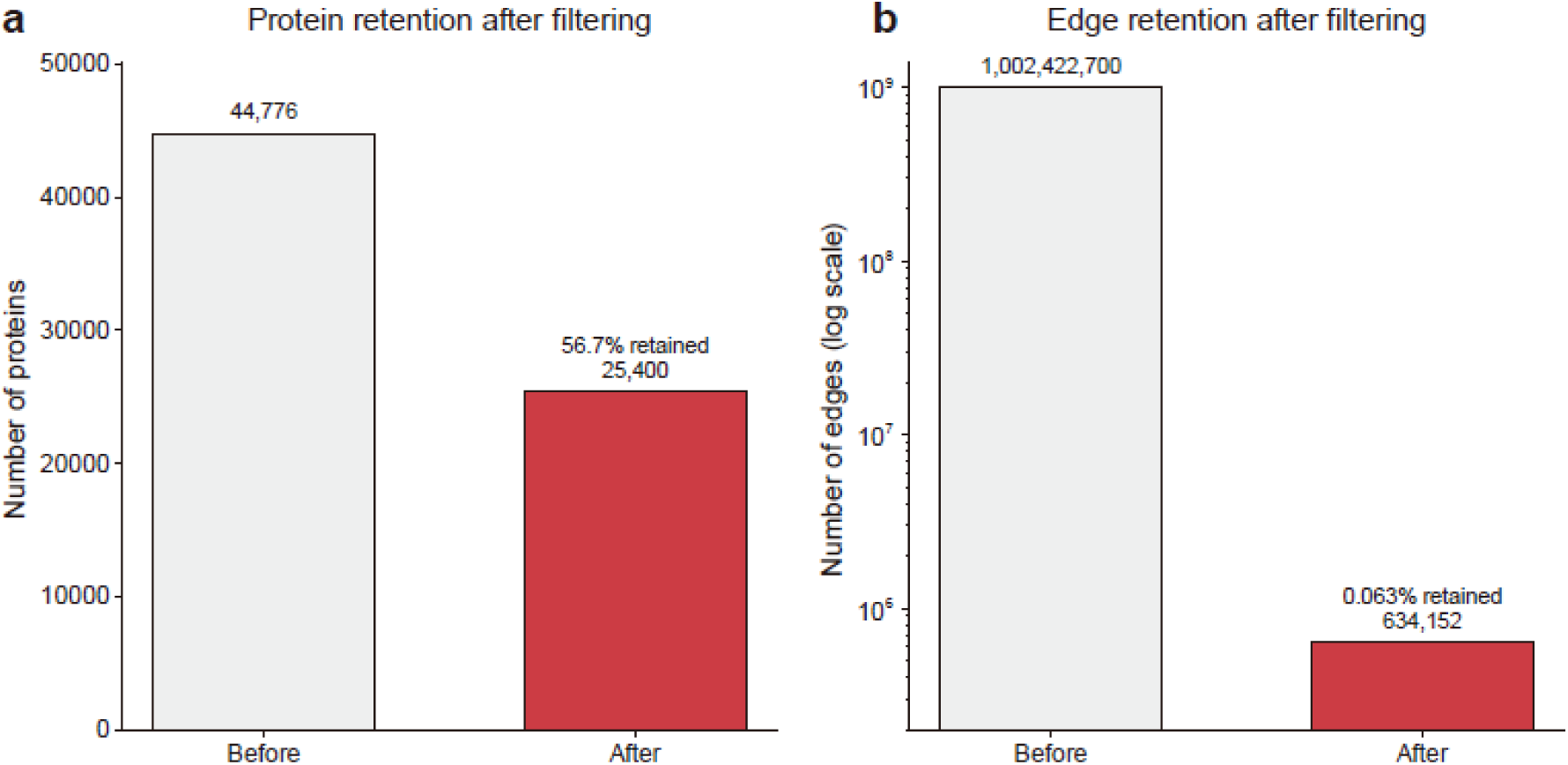
Quinoa network coverage after score thresholding. Applying the selected LoGoPPI score cutoff of 0.9 retained approximately 56.7% of quinoa proteins and 0.063% of all possible pairwise interactions, illustrating the size and sparsity of the resulting predicted interactome.

## Notes

### Competing Interest Statement

The authors have declared no competing interest.

